# Gut metabolism links precision nutrition, exercise, and healthspan in *Drosophila melanogaster*

**DOI:** 10.64898/2025.11.29.691311

**Authors:** Fangchao Wei, Shiyu Liu, Yudong Sun, Juan Liu

## Abstract

Longevity-promoting interventions commonly entail functional trade-offs, raising the unresolved question of whether lifespan extension necessarily compromises physiological performance. Here, utilizing a chemically defined diet (CDD) in *Drosophila melanogaster*, we systematically evaluated a multimodal intervention combining methionine restriction (MR), taurine supplementation (Tau), and moderate exercise. This combinatorial approach synergistically extended lifespan, preserved reproductive capacity, and improved locomotor function. Integrative targeted metabolomics and stable isotope tracing revealed increased mitochondrial tricarboxylic acid (TCA) cycle flux and enhanced redox homeostasis in the gut as central metabolic features. Notably, *Lactobacillus plantarum* was identified as a key microbial mediator responsive to dietary and behavioral stimuli, potentially coordinating host energy metabolism and maintaining physiological integrity. Together, these observations outline a “nutrition-behavior-microbiota” framework that uncouples the traditional trade-off between lifespan and functional health, offering new perspectives for promoting healthy aging.

## Introduction

A critical question in aging research is whether lifespan-extending interventions can preserve essential physiological functions, including physical performance and fertility. Methionine restriction (MR), a dietary intervention that extends lifespan across diverse species; however, often impairs reproductive capacity and functional fitness, underscoring the need to dissociate beneficial adaptations from adverse trade-offs^1,2,3,4^. Mitochondria are central mediators of dietary restriction, particularly through the regulation of redox balance and energy metabolism^4,5,6,7^. The gut acts as a critical integrator of nutritional signals, regulating systemic energy balance^8,9,10^. Accumulating evidence implicates the gut microbiota as a potent modulator of host aging and metabolism, broadly impacting physiological functions^11,12,13,14,15^. However, the mechanisms by which host-microbiota interactions influence mitochondrial function under precision dietary interventions-and how such crosstalk shapes metabolic trajectories during aging-remain poorly understood.

Taurine supplementation (Tau) and physical exercise have emerged as promising strategies to slow aging. Taurine protects mitochondria and reduces inflammation^16,17,18^, while exercise enhances metabolic fitness and preserves organ function during aging^19,20,21,22^. However, how MR, Tau, and exercise work together metabolically-and how gut bacteria might mediate their combined benefits-have not been comprehensively studied. Here, using *Drosophila melanogaster* fed a chemically defined diet (CDD)^4,7,23^, we integrate approaches from precision nutrition, microbial ecology, and metabolic flux analysis to dissect the combined effects of MR, Tau, and exercise on age-associated metabolic regulation and elucidate their coordinated modulation of host physiology through host-microbiota-mitochondria interactions.

## Results

### Synergistic enhancement of healthspan by MR and Tau in *Drosophila melanogaster*

To assess the effects of MR, Tau, and their combination (MR-Tau) on lifespan and functional aging, we designed a feeding scheme based on a CDD system in *Drosophila melanogaster* friut fly (Fig. 1a). Flies were randomized into four dietary groups: control, MR, Tau, or MR-Tau. As previously established^4^, MR contained 10% (0.402 mM) of the methionine level in the control diet and significantly extended lifespan (Extended Data Fig. 1a, b, and Fig. 1b). Tau showed a dose-dependent pro-longevity effect, with 10 mM providing maximal benefit (Extended Data Fig. 1c, d, and Fig. 1c). Both MR and Tau increased lifespan in males and females (Extended Data Fig. 1 and Fig. 1b, c), and delayed age-associated locomotor decline (Fig. 1e), consistent with previous reports^4,24,25^.

**Figure 1.**
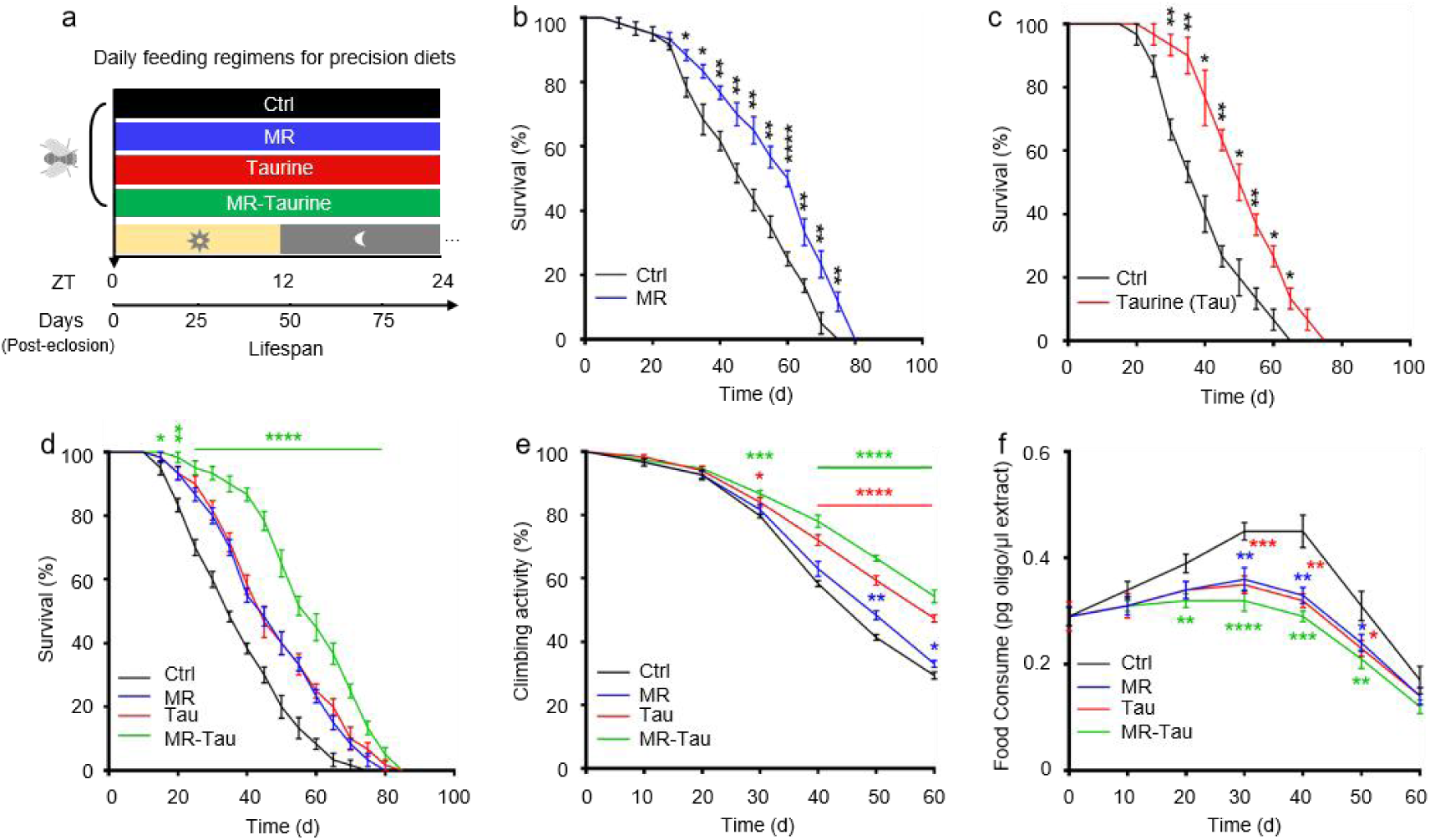
Precision methionine and taurine diet enhance healthspan in *Drosophila*. (a) Schematic of the precision diets. (b) Lifespan on MR diet (10% methionine). (c) Lifespan on taurine supplementation diet (Tau, 10mM). (d) Lifespan under different dietary conditions. (e) Climbing activity. (f) Food consumption. Lifespan (n=60), climbing activity (n=50), and food consumption (n=50) assays were performed per group, with 1:1 male-to-female ratio. All experiments were independently repeated three times. The blue, red, and green asterisks indicate the results of MR, Tau, and MR-Tau compared with the control, respectively. Data are presented as mean ± s.e.m. \**P*<0.05, \*\**P*<0.01, \*\*\**P*<0.001, \*\*\*\**P*<0.0001, *P* values were obtained by unpaired, two-tailed *t*-tests unless otherwise specified.

Methionine is metabolized through homocysteine to cysteine, a precursor for endogenous taurine biosynthesis, with these intermediates playing central roles in redox homeostasis and metabolic regulation^26^. Notably, MR-Tau intervention produced additive effects, further extending lifespan (Fig. 1d) and improving physical endurance (Fig. 1e) beyond either intervention alone.

Considering the brain’s role in regulating feeding behavior^27^, and prior evidence that both MR^28^ and taurine^29,30^ modulate hypothalamic activity to alter food intake, we evaluated feeding behavior. Indeed, flies on MR or Tau diets consumed less food, with the greatest reduction observed in the MR-Tau group (Fig. 1f). Despite reduced food intake, body weight remained comparable among groups (Extended Data Fig. 2).

Together, these results demonstrate that MR and Tau individually promote healthspan by attenuating age-related declines in survival and locomotor function. Their combination confers additive or synergistic benefits, supporting a multimodal dietary strategy that enhances healthy aging without compromising physiological integrity.

### Synergistic dietary reprogramming of systemic metabolism

To investigate how precision diets influence systemic metabolic homeostasis, we performed targeted metabolomic analysis on whole-body tissues from flies subjected to each dietary condition (Fig. 2a). Metabolite profiles were generated using liquid chromatography coupled with high-resolution mass spectrometry (LC-MS). Principal component analysis (PCA) revealed distinct metabolic profiles for the different groups (Fig. 2b), indicating that these precision diets induce broad metabolic reprogramming. Notably, the MR-Tau exhibited coordinated and pronounced changes in nucleotide, amino acid, and mitochondrial metabolites (Extended Data Fig. 3a), supporting a synergistic impact on systemic metabolism.

**Figure 2.**
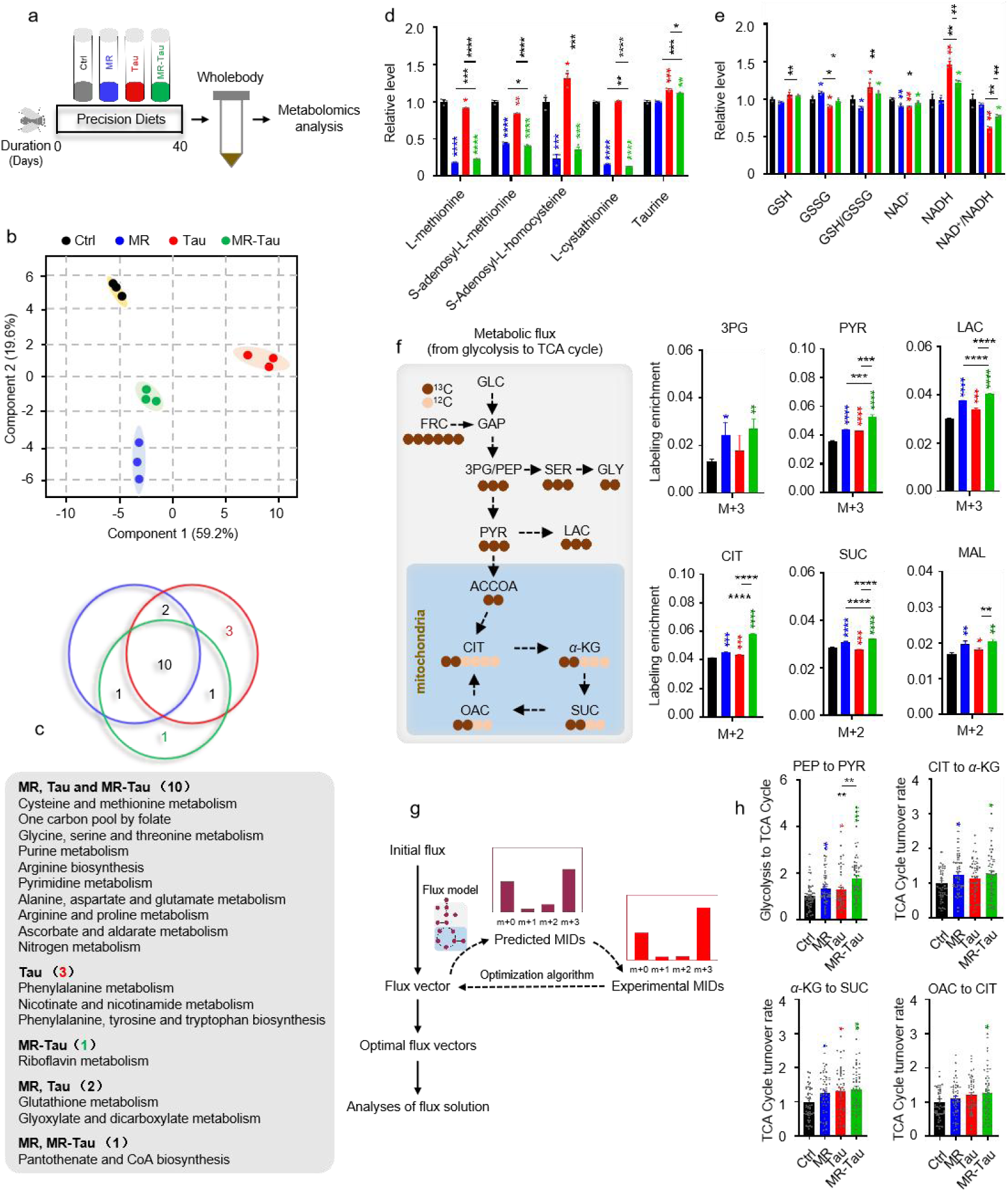
Precision diet reshapes metabolism homostasis. (a) Experimental design. (b) Principal component analysis (PCA). (c) Pathway enrichment analysis. (d) Methionine-related and taurine metabolites. (e) Redox balance. (f) Relative labeling enrichment of glycolysis-TCA cycle metabolites. (g) Central carbon metabolic flux analysis experimental design. (h) Central carbon metabolic flux analysis. n=10 flies. All experiments were independently repeated three times. Flux determination was performed by optimizing 10,000 simulated replicates. The final flux values represent the average of the 50 simulations with the smallest discrepancy in the target metabolite.The blue, red, and green asterisks indicate the results of MR, Tau, and MR-Tau compared with the control, respectively. Data are presented as mean ± s.e.m. \**P*<0.05, \*\**P*<0.01, \*\*\**P*<0.001, \*\*\*\**P*<0.0001, *P* values were obtained by unpaired, two-tailed *t*-tests unless otherwise specified. GSH, glutathione; GSSG, glutathione disulfide; NAD^+^, nicotinamide adenine dinucleotide; NADH, nicotinamide adenine dinucleotide hydrogen; MID, mass isotopomer distribution; GLC, glucose; FRC, fructose; GAP, glyceraldehyde-3-phosphate; 3PG, 3-phosphoglycerate; PEP, phosphoenolpyruvate; SER, serine; GLY, glycine; PYR, pyruvate; LAC, lactate; ACCOA, acetyl coenzyme A; CIT, citrate; *α*-KG, *α*-ketoglutarate; SUC, succinate; MAL, malate; OAC, oxaloacetate.

Pathway enrichment analysis identified ten pathways commonly altered across all interventions, including one-carbon metabolism (methionine and cysteine metabolism, folate cycle), amino acid metabolism (glycine, serine, and threonine metabolism; arginine and proline metabolism), and purine metabolism (Fig. 2c). Consistently, the quantification of methionine- and taurine-related metabolites from these pathways revealed significant changes across interventions (Fig. 2d). Remarkably, MR-Tau uniquely affected pathways related to glutathione biosynthesis and mitochondrial redox regulation (Fig. 2e), indicating altered antioxidant capacity and mitochondrial function.

Carnitine, essential for fatty acid import into mitochondria, serves as a key indicator of TCA cycle activity and energy flux^31,32,33^. Age-associated declines in carnitine metabolism impair mitochondrial function and lipid utilization^34,35^, while supplementation improves energy metabolism and insulin sensitivity in aged individuals^35^. A marked accumulation of carnitine species was observed under MR-Tau (Extended Data Fig. 3b) and upregulated endogenous carnitine biosynthesis pathways (Fig. 2c), consistent with increased mitochondrial fatty acid transport and *β*-oxidation, which may contribute to maintaining metabolic integrity during aging.

To further assess the effects of different dietary conditions on mitochondrial function and redox balance, we performed stable isotope tracing. This analysis revealed that MR-Tau markedly enhanced TCA cycle flux (Fig. 2f, g, h), accompanied by increased production of one-carbon donors and improved redox homeostasis-metabolic hallmarks of resilience that support healthy aging.

### MR-Tau intervention modulates gut mitochondrial metabolism and supports systemic metabolic homeostasis

The MR-Tau combination significantly reduced food intake (Fig. 1f) without affecting body weight (Extended Data Fig. 2). Given the central regulation of feeding behavior^27,36^, we first examined the impact of precision diets on brain metabolism. Metabolomic profiling of fly heads revealed distinct signatures across dietary conditions (Extended Data Fig. 4a, b), with MR generally associated with reduced levels of brain metabolites, whereas Tau or MR-Tau tended to preserve or elevate these levels (Extended Data Fig. 4c). MR markedly depleted methionine-related metabolites, whereas taurine supplementation elevated methionine cycle intermediates and partially restored MR-induced reductions (Extended Data Fig. 4d). Notably, MR-Tau markedly increased whole-body mitochondrial TCA flux (Fig. 2f, h) and elevated the pyruvate/lactate ratio in heads (Extended Data Fig. 4e), indicating enhanced systemic oxidative metabolism and a potential synergistic effect on brain energy metabolism and feeding regulation.

Given the gut’s pivotal role in nutrient sensing^8,9^ and its close association with host metabolism and the microbiota^13,14^, we next evaluated how MR, Tau, and MR-Tau influence gut metabolic homeostasis. Following established protocols^37^, we focused on the midgut (Fig. 3a), the primary site of nutrient absorption and functionally analogous to the mammalian small intestine^38^. Beyond digestion, the midgut contributes to immunity and systemic metabolism via dynamic host-microbe interactions^37,38,39^. Targeted metabolomic profiling revealed distinct dietary signatures (Extended Data Fig. 5a), with remodeling in nucleotides, amino acids, and intermediary metabolites (Extended Data Fig. 5b). Pathways associated with methionine and taurine metabolism (Fig. 3b and Extended Data Fig. 5c), redox homeostasis (Fig. 3c), and carnitine metabolism (Extended Data Fig. 5d) were notably changed, with the most pronounced alterations in the MR-Tau group. These changes align with previous findings linking improved antioxidant capacity and mitochondrial function to increased longevity^4,5,6,40^.

**Figure 3.**
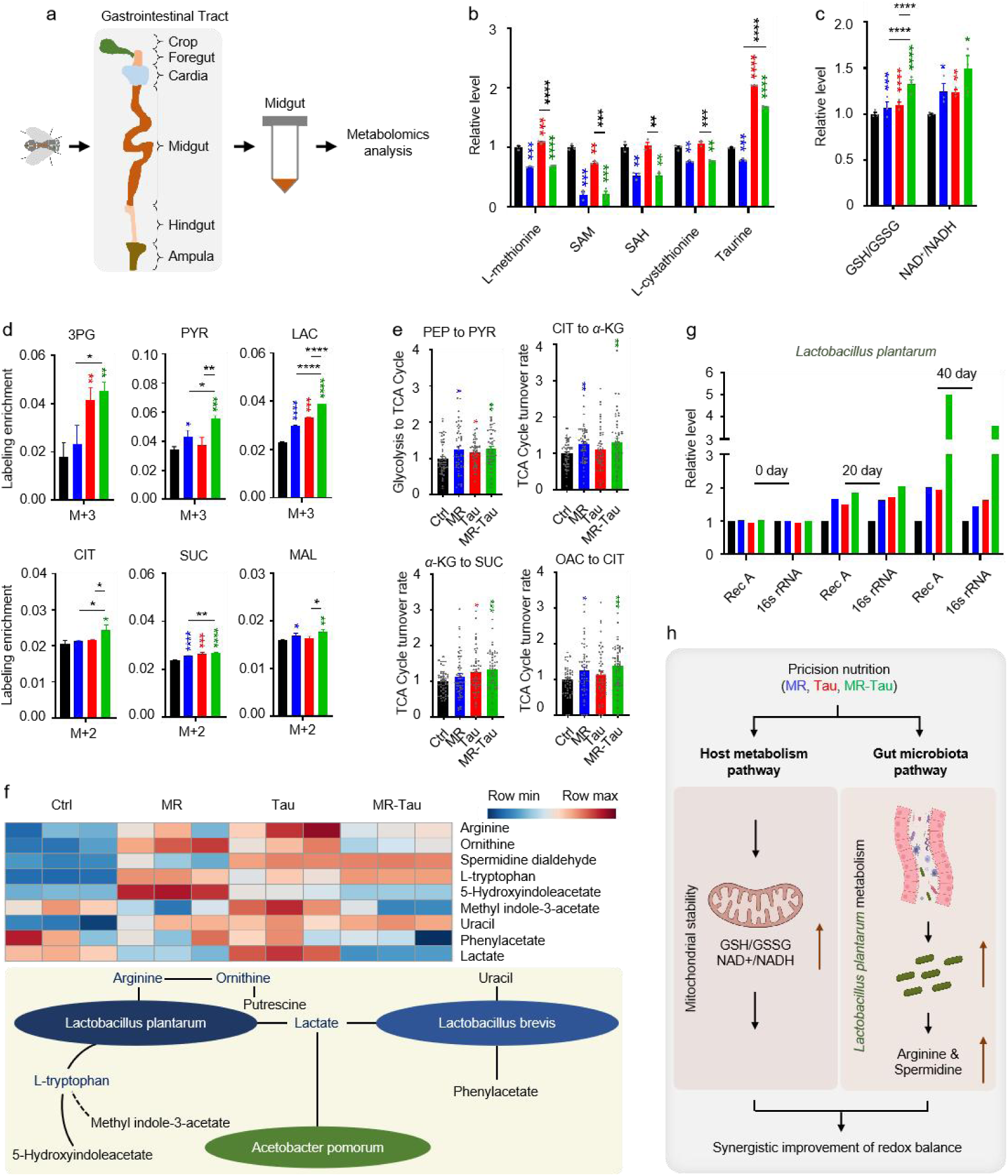
Precision nutrition regulates gut and microbial metabolic homeostasis. (a) Experimental design. (b) Methionine-related and taurine metabolites. (c) Redox balance. (d) Relative labeling enrichment of glycolysis-TCA cycle metabolites. (e) Central carbon metabolic flux analysis. (f) Metabolic network of gut microbiota. (g) Effect of precision diet on gut *L. plantarum* abundance. (h) Model. n=30 flies. All experiments were independently repeated three times (h, all reactions were run in five technical replicates). Flux determination was performed by optimizing 10,000 simulated replicates. The final flux values represent the average of the 50 simulations with the smallest discrepancy in the target metabolite. The blue, red, and green asterisks indicate the results of MR, Tau, and MR-Tau compared with the control, respectively. Data are presented as mean ± s.e.m. \**P*<0.05, \*\**P*<0.01, \*\*\**P*<0.001, \*\*\*\**P*<0.0001, *P* values were obtained by unpaired, two-tailed *t*-tests unless otherwise specified. SAM, S-adenosyl-L-methionine; SAH, S-adenosyl-L-homocysteine.

TCA cycle flux was increased in the midguts of MR-Tau flies (Fig. 3d, e), indicative of enhanced mitochondrial function. Pathway enrichment analysis further identified eight metabolic pathways consistently altered across all interventions (Extended Data Fig. 5c), including one-carbon metabolism and multiple amino acid-related pathways. Taken together, these results indicate that MR-Tau is associated with enhanced gut mitochondrial activity and one-carbon metabolism, which may contribute to improved redox balance and systemic metabolic stability-features often linked to healthy aging.

### MR-Tau enhances microbial metabolic activity and supports gut homeostasis

Gut microbiota are integral to host metabolic regulation, immune function, and redox balance, thereby influencing healthspan and aging^15,41,42,43,44,45,46^. Among the evolutionarily conserved commensals shared between *Drosophila* and mammals are *Lactobacillus plantarum* (*L. plantarum*) and *Lactobacillus brevis* (*L. brevis*), which serve functionally analogous roles across species^13,14,15,37,43^. In contrast, *Acetobacter pomorum* (*A. pomorum*), a dominant fly-specific symbiont, is well characterized for its effects on host development and metabolic regulation^43^. Several microbial taxa are linked to specific metabolic pathways and their associated metabolites: *L. plantarum* is involved in amino acid catabolism and serotonin/indole signaling pathways, producing metabolites such as arginine^47,48^, tryptophan^49,50^, and indole derivatives^51,52^; *L. brevis*^44,53,54^ participates in nucleotide and aromatic amino acid metabolism, generating uracil and phenylacetic acid; and *A. pomorum* contributes to lactate catabolism, which may modulate host energy balance through central carbon metabolism^12^.

To assess whether dietary interventions influence microbiota-derived metabolites, we profiled representative compounds. Taurine supplementation was associated with increased levels of arginine, ornithine, and tryptophan, as well as an enhancement of spermidine metabolism (Fig. 3f), consistent with elevated microbial metabolic activity and potentially strengthened host-microbe interactions^47,48^. These findings align with evidence that tryptophan-enriched diets promote *Lactobacillus*-linked longevity benefits^50^.

We next quantified bacterial abundance using conserved molecular markers (*recA* and *16S rRNA*)^13,55,56,57,58^. Longitudinal analysis revealed a gradual increase in *L. plantarum* abundance, with the most pronounced change in the MR-Tau group (Fig. 3g). *A. pomorum* also showed a moderate rise (Extended Data Fig. 6b), while *L. brevis* remained unchanged (Extended Data Fig. 6a). These observations suggest that *L. plantarum* could contribute to the metabolic effects of MR-Tau, potentially via arginine biosynthesis and support of host energy and redox balance. Altogether, these results indicate that precision dietary interventions can modulate the gut microbial ecosystem, with MR-Tau associated with enhanced *L. plantarum* representation and microbial metabolic activity. Such changes may help maintain intestinal functional stability and support systemic metabolic homeostasis, features relevant to healthy aging (Fig. 3h).

### MR-Tau combined with exercise supports lifespan extension, physical performance, and metabolic health

Exercise is a well-established modulator of systemic metabolism and a protective factor against age-related functional decline^19,20,21,22^ but its interactions with precision dietary strategies remain insufficiently understood. Building on the metabolic and microbial improvements associated with MR-Tau, we next investigated whether combining this dietary approach with moderate physical activity could produce additive or synergistic effects on healthspan. To address this, we implemented a moderate-intensity exercise regimen across different dietary conditions (Fig. 4a). Exercise extended lifespan in all groups, with the most pronounced longevity observed in the MR-Tau-exercise cohort (Fig. 4b), indicating a possible synergistic interaction.

**Figure 4.**
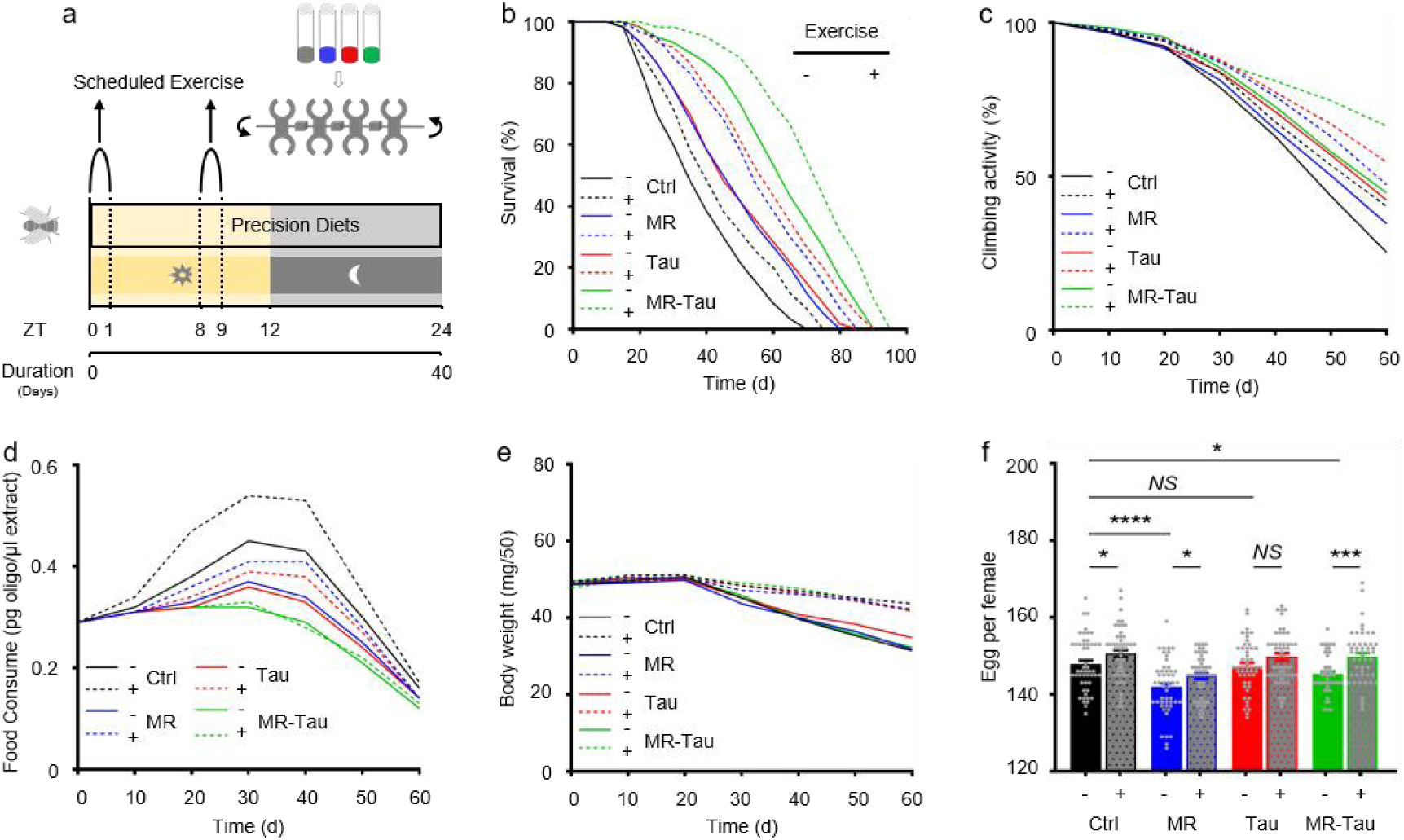
Exercise synergizes with precision nutrition-enhanced anti-aging effects. (a) Experimental design. (b) Lifespan. (c) Climbing activity. (d) Food consumption. (e) Body weight. (f) Egg production. Lifespan (n=60), climbing (n=50), food consumption (n=50), body weight (n=150) and egg production (n=50) assays were performed per group, with 1:1 male-to-female ratio. All experiments were independently repeated three times. Data are presented as mean ± s.e.m. *NS*, not significant *P*>0.05, \**P*<0.05, \*\*\**P*<0.001, \*\*\*\**P*<0.0001, *P* values were obtained by unpaired, two-tailed *t*-tests unless otherwise specified.

In parallel, longitudinal assessments of locomotor function indicated that all interventions attenuated age-related decline, with MR-Tau-exercise maintaining the highest average physical performance over time (Fig. 4c). These findings are consistent with improved neuromuscular preservation under combined intervention^21^. Interestingly, although exercise generally increased food intake, MR-Tau-fed flies sustained lower feeding rates even during physical activity (Fig. 4d), indicating that diet composition remained a dominant driver of feeding behavior. Body weight was stable across conditions (Fig. 4e).

Notably, exercise also partially mitigated the reduced fecundity observed with MR and MR-Tau (Fig. 4f), aligning with previous evidence that nutritional and behavioral interventions can jointly influence reproductive and metabolic outcomes^59^. Taken together, these results indicate that combining MR-Tau with moderate exercise confers broad physiological benefits-including extended lifespan, maintained locomotor capacity, metabolic stability, and partial preservation of reproductive function.

### MR-Tau-exercise remodels the gut metabolic microenvironment

The gut is a central regulator of systemic energy metabolism and redox homeostasis^57^. Consistent with prior reports^58,59^, dietary composition modulated the gut microenvironment through changes in microbial composition and metabolic function (Fig. 3f, g, Extended Data Fig. 5 and Extended Data Fig. 6). To investigate how MR-Tau-exercise shapes host-microbe metabolic crosstalk, we assessed redox balance, lipid metabolism, microbial dynamics, and central metabolic fluxes. Redox-sensitive metabolites increased stepwise from control to MR-Tau, peaking in the MR-Tau-exercise group (Fig. 5a), consistent with enhanced redox buffering capacity and mitochondrial function^4^. This was accompanied by increased expression of *L. plantarum recA* and *16S rRNA* (Fig. 5e), while *L. brevis* remained unchanged (Extended Data Fig. 7a), suggesting a species-specific microbial response.

**Figure 5.**
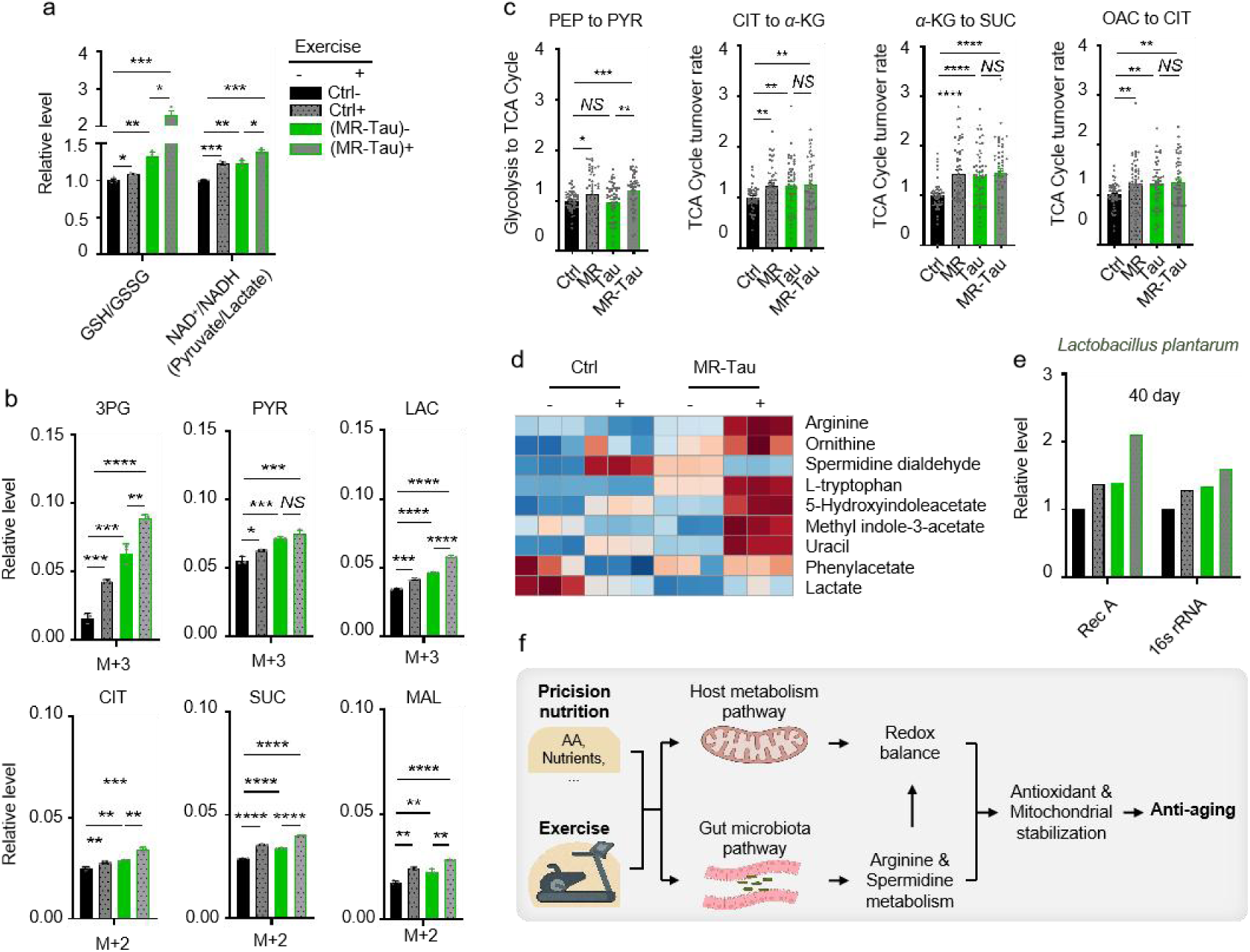
Exercise synergizes with precision nutrition to remodel gut metabolism and reinforce anti-aging capacity. (a) Redox balance. (b) Relative labeling enrichment of glycolysis-TCA cycle metabolites. (c) Central carbon metabolic flux analysis. (d) Metabolites linked to gut bacteria. (e) Effect of combined intervention on *L. plantarum* abundance. (f) Model. n=30. All experiments were independently repeated three times (e, all reactions were run in five technical replicates). Flux determination was performed by optimizing 10,000 simulated replicates. The final flux values represent the average of the 50 simulations with the smallest discrepancy in the target metabolite. Data are presented as mean ± s.e.m. *NS*, not significant; *P*>0.05, \**P*<0.05, \*\**P*<0.01, \*\*\**P*<0.001, \*\*\*\**P*<0.0001, *P* values were obtained by unpaired, two-tailed *t*-tests unless otherwise specified.

Consistent with elevated mitochondrial lipid utilization, free, short-, and medium-chain carnitine levels were increased in MR-Tau and remained above baseline with exercise (Extended Data Fig. 7c). Stable isotope tracing analysis further indicated that MR-Tau-exercise produced the strongest activation of central carbon metabolism among all tested conditions (Fig. 5b, c and Extended Data Fig. 8c), supporting synergistic engagement of mitochondrial energy pathways.

Microbial-derived metabolites, including *L. plantarum*-associated arginine and tryptophan derivatives, were significantly enriched under MR-Tau-exercise (Fig. 5d), consistent with enhanced microbial biosynthetic output. Interestingly, *A. pomorum* abundance was not altered by exercise alone, but was increased in MR-Tau and remained elevated when exercise was added (Extended Data Fig. 7b), suggesting that diet exerts the primary influence on microbial composition, while exercise may augment diet-induced changes.

These alterations were consistently observed across whole-body, gut, and brain metabolomes (Extended Data Fig. 8), indicating that MR-Tau-exercise integrates host and microbial metabolic responses across multiple tissues. Overall, these findings indicate that this combined intervention reconfigures the gut metabolic niche-enhancing redox capacity, supporting lipid utilization, and modulating microbial function-in a manner that aligns with improved metabolic homeostasis and functional health.

## Discussion

Our study demonstrates that MR-Tau-exercise robustly extends lifespan in *Drosophila melanogaster* without compromising physiological integrity. This multimodal approach enhances mitochondrial redox balance, energy metabolism, and gut microbial composition-particularly enriching *L. plantarum*-which together support systemic metabolic resilience (Fig. 5f).

While dietary interventions like MR or caloric restriction have been widely reported to promote longevity^1,2,4,7,60^, these approaches often involve trade-offs, including reduced fertility and physical performance. In contrast, the MR-Tau-exercise combination not only prolongs lifespan but also preserves locomotor function, metabolic homeostasis, and reproductive output. Mechanistically, this intervention enhances gut antioxidant capacity (Fig. 5a), TCA cycle flux (Fig. 5b, c), and *L. plantarum*-derived metabolites (e.g., arginine and tryptophan derivatives), which may contribute to improved host mitochondrial function via metabolite-mediated signaling pathways. Although these associations are consistent with previous work implicating *Lactobacillus* species in healthspan regulation^61^, establishing causality will require further studies using germ-free models, microbial transplantation, or targeted microbial perturbations. Together, these findings support a precision aging framework that integrates dietary, behavioral, and microbiota-based strategies.

Despite compelling evidence from this *Drosophila* model, future investigations in mammalian systems are essential to assess translational potential. Notably, MR has demonstrated lifespan- and healthspan-extending effects in rodents^7^, and taurine supplementation has recently been associated with improved metabolic health and delayed aging phenotypes in mice and non-human primates^18^. Additionally, exercise is a well-established modulator of mitochondrial function and systemic metabolism across species^20,21,22^. These independent lines of evidence suggest that the synergistic benefits observed in flies might extend to mammals. However, rigorous validation incorporating tissue-specific metabolomics, microbiome profiling, and behavioral assessments in murine models will be critical to evaluate efficacy, safety, and tissue-specific effects within more complex organisms, providing foundational data for the clinical translation of precision aging strategies tailored to individual metabolic and microbial states.

Tissue-specific metabolomic profiling revealed coordinated inter-organ remodeling under MR-Tau-exercise. In the brain, TCA cycle activity and neuroactive metabolite levels were restored (Extended Data Fig. 4c), while in the gut, energy and redox pathways were upregulated (Fig. 3c, d, e and Extended Data Fig. 5), indicating coordinated metabolic responses across these tissues. qPCR analysis further confirmed the enrichment of *L. plantarum* under the intervention (Fig. 3g, Fig. 5e), accompanied by shifts in associated metabolites, which correlated with host metabolic improvements (Fig. 4). Together with prior evidence^47,48,49,50^ linking *L. plantarum* to amino acid metabolism, redox balance, and longevity, these results highlight a biologically plausible contribution of microbiota remodeling to the observed neuro-metabolic benefits. Mechanistic studies integrating behavioral assays, inter-organ flux tracing, and microbial genetics will be essential to dissect how this “diet-exercise-microbiota” axis may shape gut-brain communication and systemic resilience during aging.

Whether taurine functions as an aging biomarker remains context dependence^16,17,18,62^. In our study, taurine conferred robust anti-aging effects, particularly when combined with MR and exercise, acting as a conditional longevity modulator and underscoring the value of synergistic nutritional interventions. Collectively, these findings reinforce a precision aging framework that integrates dietary, behavioral, and microbiota-based strategies. This tri-modal approach represents a promising avenue to delay age-related functional decline while maintaining systemic resilience. Future mammalian studies will be essential to translate these insights into clinically relevant interventions tailored to individual metabolic and microbial states.

## Methods

### Fly Stocks and Maintenance

*Drosophila melanogaster* (*w^11^*^18^) flies were obtained from the Bloomington Stock Center and maintained at 25 °C with 60% humidity under a 12:12 h light-dark cycle. Newly eclosed flies (≤8 h) were collected, grouped (25 per vial), and transferred to fresh food every 2-3 days.

### Diet

Flies were maintained on a chemically defined diet (CDD) adapted from published protocols^4,23,63,64,65^. Modifications to the CDD included changes in agar type, sucrose concentration, nucleoside composition, and amino acid ratios to reflect the *Drosophila* exome^4,23,65^. The full formulation of the CDD is provided in Supplementary Data 1. Variants of the CDD were prepared for specific experimental needs, such as methionine-restricted, nutrient-enriched, or isotope-labeled diets (U-^13^C_6_-Sucrose (fructose), Cambridge, Cat. CLM-9811-PK), with the fructose labeling rate shown in Supplementary Data 2. Diets were stored at 4 °C. For reproductive assays, experimental diets were provided to one sex while the other remained on control food.

### Lifespan Assay

Flies were collected within 8 hours post-eclosion, sorted under CO₂, and randomly allocated to different diets (each group contained three replicates with 10 flies per vial). They were transferred to fresh food every 2-3 days, and deaths were recorded daily. All flies used in lifespan analysis were kept virgin to exclude mating-related effects.

### Food Intake Assay with DNA Oligomer Incorporation and qPCR

Three DNA oligomers were prepared in ddH₂O at final concentrations of 1.75 µg/µL, 2.5 µg/µL and 3.5 µg/µL, respectively, based on previous studies^66,67^. A total of 90 µL of the mixture was applied evenly onto the surface of fly food and air-dried. Sex-separated flies were transferred to the food-containing vials and maintained at 25 °C overnight. The following day, flies were briefly anesthetized with CO₂, and 10 flies per sample were collected for analysis.

For DNA barcode qPCR, flies were homogenized in 70 µL squishing buffer (10 mM Tris-HCl pH 8.2, 25 mM NaCl, 1 mM EDTA, 200 µg/mL proteinase K) using 0.5 mm zirconium beads. Homogenates were incubated at 37 °C for 40 min, followed by heat inactivation at 95 °C for 5 min. Samples were centrifuged at 10,000g for 10 min, and 10 µL of the supernatant was used for qPCR using Power SYBR™ Green PCR Master Mix (Thermo Fisher, Cat. 4367659) on a Viia 7 Real-Time PCR System (Roche). Amplification was performed at 60 °C for 40 cycles.

The sequences of the DNA oligomers and qPCR primers used are as follows: DNA Oligomer 1:

5’-ACCTACACGCTGCGCAACCGAGTCATGCCAATATAAGCAGATTAGCATTACTTTGAGCAACGTATCGGCGATCAGTTCGCCAGCAGTTGTAATGAGCCCC-3’

Forward qPCR Primer 1:5’-GCAACCGAGTCATGCCAATA-3’ Reverse qPCR Primer 1:5’-TTACAACTGCTGGCGAACTG-3’

DNA Oligomer 2:

5’-GGGCAGCAGGATAACTCGAATGTCTTAGTGCTAGAGGCTTGGGGCGTGTAAGTGTATCGAAGAAGTTCGTGTTAAACGCTTTGGAATGACTGTAATGTAG-3’

Forward qPCR Primer 2:5’-CAGCAGGATAACTCGAATGTCTTA-3’ Reverse qPCR Primer 2:5’-CAGTCATTCCAAAGCGTTTAACA-3’

DNA Oligomer 3:

5’-CTGTAGTATTCGTCCGACGTTCCTCTCTGCTCGGGTACGCGACGAAGGCTCTACTGGCAGTCGAGATTATCGTACAATTTAGTTCGGCCAACCTGAAGCT-3’

Forward qPCR Primer 3:5’-CTGTAGTATTCGTCCGACGT-3’ Reverse qPCR Primer 3:5’-AGCTTCAGGTTGGCCGAACT-3’

### Egg Counting

To assess reproductive output, mated females were transferred to fresh diet vials daily. Egg production was quantified from day 3 post-mating over a 30-day period. For reproductive assays, only one sex (male or female) was exposed to the experimental diet, while the mating partner remained on control food. Mating was conducted for 5 hours on control media^4,68^. Flies used for egg counting were allowed to mate only once.

### Climbing Assay and Body Weight Measurement

Climbing ability was assessed as previously described^4,69^. Briefly, 10 single-sex flies were transferred into a 23 × 95 mm empty vial, tapped down three times, and the number of flies reaching the top within 20 seconds was recorded. Three vials per group were tested under each condition. Tests were performed every 10 days over a 2-month period.

For body weight, 50 flies were collected, weighed every 10 days, and followed up for 2 months. Each experiment was performed on at least 150 flies (3 vials of 50 flies per condition) and repeated 3 times.

### *Drosophila* exercise system

Based on previous studies^70^, a TreadWheel apparatus was used for exercise training in *Drosophila*. Each device accommodates 12 fly vials mounted on four axles and fits within a standard *Drosophila* incubator. Vials were rotated lengthwise at a constant speed of 5 revolutions per minute (RPM) by an electric motor, continuously altering the vertical orientation of the vial. This gentle, sustained rotation exploits the flies’ innate negative geotaxis, providing a consistent climbing stimulus throughout the training period.

### Tissue Preparation

Flies were anesthetized with CO₂ first. Whole-body samples were prepared using 10 flies per group, snap-frozen in liquid nitrogen before storage at - 80 °C.

For head samples, heads were rapidly dissected under a stereomicroscope using a scalpel. Dissected heads were immediately transferred into Eppendorf tubes and snap-frozen in liquid nitrogen before storage at -80 °C. Each head sample group consisted of heads from 30 flies.

For midgut samples, the digestive system was dissected following established^39,71^. The midgut was isolated, quickly transferred into Eppendorf tubes, snap-frozen in liquid nitrogen, and stored at -80 °C. Each midgut sample group consisted of midguts from 30 flies.

### RNA Extraction and qPCR Analysis

Total RNA was extracted from dissected *Drosophila* midguts using the (QIAwave RNA Mini Kit, Cat. 74536) following established protocols^72^. To remove contaminating genomic DNA, RNA samples were treated with RQ1 RNase-Free DNase (Promega, Cat. M6101). cDNA was synthesized from total RNA using the iScript cDNA synthesis kit (Bio-Rad, Cat. 170-8891).

Quantitative PCR (qPCR) was performed using Power SYBR ™ Green PCR Master Mix (Thermo Fisher, Cat. 4367659) on a Viia 7 Real-Time PCR System (Roche). We focused on three representative bacterial species commonly found in the *Drosophila* midgut: *Lactobacillus plantarum*, *Lactobacillus brevis*, and *Acetobacter pomorum*.

Gene-specific primers targeting *16S rRNA*, *recA*, and *aldo/keto reductase* (AKR, *L. brevis*-specific) genes were designed based on published sequences^73,74,75,76,77^. Total bacterial 16S rRNA served as an internal control for normalization^78^. Primer sequences are listed below:

### Lactobacillus plantarum

*16S rRNA*^73^: Lp-16S-Forward 5’-GTGSTGCAYGGYTGTCGTCA-3’; Lp-16S-Reverse 5’-ACGTCRTCCMCACCTTCCTC-3’ *recA*^74^: Lp-recA-Forward 5’-CCGTTTCTGCGGAACACCTA-3’; Lp-recA-Reverse 5’-TCGGGATTACCAAACATCAC-3’

### Lactobacillus brevis

*16S rRNA*^75^: Lb-Forward 5’-ATTTTGTTTGAAAGGTGGCTTCGG-3’; Lb-Reverse 5’-ACCCTTGAACAGTTACTCTCAAAGG-3’

*AKR*^76^: Lb-recA-Forward 5’-AATTGATTTTCATACCGCAGAA-3’; Lb-recA-Reverse 5’-TTGGCACCGCATGATGTG-3’

### Acetobacter pomorum

*16S rRNA*^77^: Ap-16S-Forward 5’-TCAAGTCCTCATGGCCCTTATG-3’; Ap-16S-Reverse 5’-TCGAGTTGCAGAGTGCAATCC-3’

### Total bacterial

*16s rRNA*^78^: 341F-5’-ACTCCTACGGGAGGCAGCAG-3’; 534R-5’-ATTACCGCGGCTGCTGG-3’

The amplification protocol was as follows: initial denaturation at 95 °C for 3 min, followed by 40 cycles of 95 °C for 15 s, 55 °C for 30 s, and 72 °C for 30 s. Melting curve analysis at the end of the run to confirm primer specificity. Each reaction was performed with five technical replicates. Relative abundance of bacterial species was calculated using the *ΔCt* method, normalized to total bacterial *16S rRNA* levels.

### Metabolite Extraction

10 flies or 30 flies’ heads or midguts were collected under light CO₂ anesthesia, snap-frozen in liquid nitrogen, and stored at -80 °C. Tissues were ground using a CryoMill, and metabolites were extracted as previously described^79^. Supernatants were dried in a vacuum concentrator at room temperature and reconstituted in 30 μl of solvent (15 μl water followed by 15 μl methanol/acetonitrile, 1:1, v/v). A 3 μl aliquot was analyzed by liquid chromatography coupled with high-resolution mass spectrometry using water with 5 mM ammonium acetate (pH 6.9) as mobile phase A and 100% acetonitrile as mobile phase B.

Metabolite separation and detection were performed using an Ultimate 3000 UHPLC system (Dionex) coupled to a Q Exactive mass spectrometer (Thermo Scientific). Metabolites were separated at room temperature using a hydrophilic interaction chromatography method (HILIC) method with an XBridge Amide column (100 × 2.1 mm, 3.5 μm; Waters). Mobile phase composition and gradient conditions were described previously^79^. Briefly, the linear gradient of mobile phase B was: 0 min, 85%; 1.5 min, 85%; 5.5 min, 35%; 10 min, 35%; 10.5 min, 35%; 10.6 min, 10%; 12.5 min, 10%; 13.5 min, 85%; and 20 min, 85%. The flow rate was 0.15 ml/min (0-5.5 min), 0.17 ml/min (6.9-10.5 min), 0.3 ml/min (10.6-17.9 min), and 0.15 ml/min (18-20 min). The mass spectrometer was equipped with a heated electrospray ionization (HESI) source. Key parameters were: evaporation temperature, 120 °C; sheath gas, 30; auxiliary gas, 10; sweep gas, 3; spray voltage, 3.6 kV (positive mode) or 2.5 kV (negative mode); capillary temperature, 320 °C; and S-lens level, 55. Full MS scans were acquired over m/z 70-900 at 70,000 resolution, with a maximum injection time of 200 ms and automatic gain control (AGC) target of 3 × 10⁶ ions. Customized mass calibration was performed before data acquisition.

### Metabolomics Data Processing and Metabolic Flux Analysis (MFA)

LC-MS peak extraction and integration were performed using Sieve 2.2 software (Thermo Scientific). Integrated peak intensities were used for downstream analysis. Natural isotope abundance correction was applied as previously described^80^ for isotope tracing studies.

A metabolic network model was constructed covering glycolysis, the TCA cycle, the pentose phosphate pathway, one-carbon metabolism, and amino acid biosynthesis^4,81^. To minimize random errors from batch effects, mass isotopomer distributions (MIDs) of metabolites were averaged across biological replicates. The average MID of each metabolite was predicted as the weighted sum of its precursor MIDs, based on presumed generating fluxes. These average MID data are fitted with the following procedure: MIDs of all target metabolites are predicted by averaging the MID of the precursors, weighted by the corresponding generating fluxes with the presumed value:

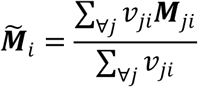

*M_i_*: predicted MID vector of metabolite *i*; *M_ji_*: MID of metabolite *i* produced from a substrate *j*; *v_ji_*: the flux from *j* to *i*.

If *M_ji_* is still unknown, it can be deduced by the same procedure until MIDs of all precursors are known. Then, the difference between the predicted and experimental MIDs of target metabolites, was evaluated by sum of squared error:

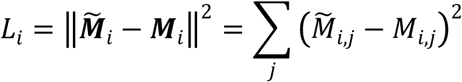

*L_i_*: difference of target metabolite *i*.*M_i,j_*, *M_i,j_*: element *j* in vector *M_i_* and *M_i_* Sum of *L_i_* for all target metabolites was regarded as the total difference *L*_total_ to minimize by adjusting flux vector *v* = {*v_i_*} that including all fluxes. Therefore, an optimization problem was defined as:

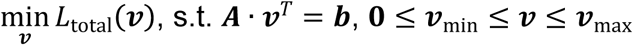

*A v^T^* = *b*: flux balance requirement and other equality constraints; *v*_min_, *v*_max_ lower and upper bounds for composite vector *v* . The solution of this optimization problem *v*^∗^ gives a combination of all fluxes in the network model that fits MID data. Method utilized to solve this optimization problem refers to previous publication^82,83^. The optimization was repeated 10,000 times, and the 50 solutions with the lowest final *L*_total_ were selected as the final solution set to ensure precision, accuracy, and robustness. In line with our previous study using the model^4^, the method has been further applied and systematically optimized in this study.

### Statistics and Reproducibility

All experiments were performed with at least three biological replicates and repeated independently two to three times. Data are presented as mean ± s.e.m. unless otherwise indicated. *P* values were calculated using two-tailed Student’s t-tests unless stated otherwise. Sample sizes were not pre-determined statistically but were consistent with those in previous studies^1,4^. Data distribution was assumed to be normal, though not formally tested. In *Drosophila* phenotype experiments, the investigators were not blinded to allocation during the experiments and outcome assessment due to the nature of the experimental setup and practical limitations, which is in line with standard practices in the field. In mass spectrometry experiments, data collection and analysis were performed in a blind manner, and steps were taken to avoid batch effects.

## Data Availability

All data supporting the findings of this study are available at https://github.com/LocasaleLab/MR_Tau_Ex_2025. Source data are provided in this paper. Metabolite data analysis was performed using MetaboAnalyst 6.0 (http://www.metaboanalyst.ca/ MetaboAnalyst/) and GENE-E (https://software.broadinstitute.org/GENE-E/) with pathway annotations from the Kyoto Encyclopedia of Genes and Genomes (KEGG) (http://www.genome.jp/kegg/).

## Code Availability

All scripts were implemented in Python 3.8. Source code is available at https://github.com/cmplab-cimr/202508_Fangchao_MR_Tau_Exercise.

## Acknowledgments

We thank Dr. Jason W. Locasale for supporting this work through the National Institutes of Health grant R01CA193256 (principal investigator) and for his insightful suggestions. This work was further supported by departmental funds from the Duke Department of Pharmacology and Cancer Biology. We regret that some relevant work could not be cited due to space limitations. Schematics of the *Drosophila* model, mitochondria, and midgut were created with *BioRender.com*.

## Author Contributions Statement

F.C.W. conceived and designed the study, performed experiments, analyzed data, and drafted the manuscript. S.Y.L. developed the metabolic flux calculation method. F.C.W and Y.D.S. optimized the chemically defined diet.

Y.D.S. and J.L. assisted in metabolomics data analysis. F.C.W. supervised the project. All authors reviewed and approved the final manuscript.

## Competing Interests Statement

The authors declare no competing interests.

---

**Extended Data Figure 1.**
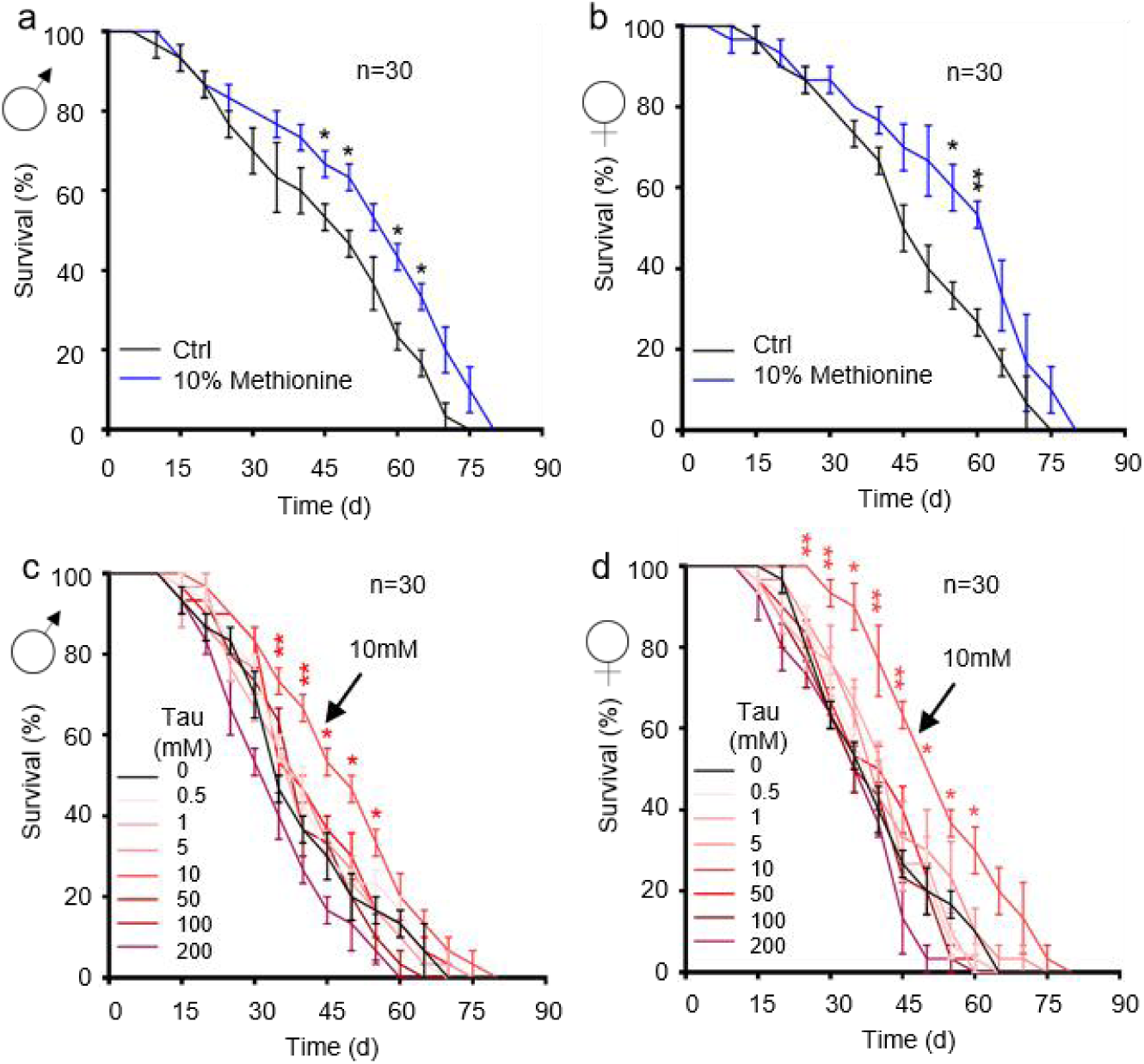
Methionine restriction (MR) and taurine supplementation (Tau) enhance healthspan in *Drosophila*. (a-b) Lifespan on MR (10% methionine); male (a), female (b). (c-d) Lifespan on Tau; male (c), female (d). Lifespan (n=60) assays were performed per group, with 1:1 male-to-female ratio. All experiments were independently repeated three times. Data are presented as mean ± s.e.m. \**P*<0.05, \*\**P*<0.01, *P* values were obtained by unpaired, two-tailed *t*-tests unless otherwise specified.

**Extended Data Figure 2.**
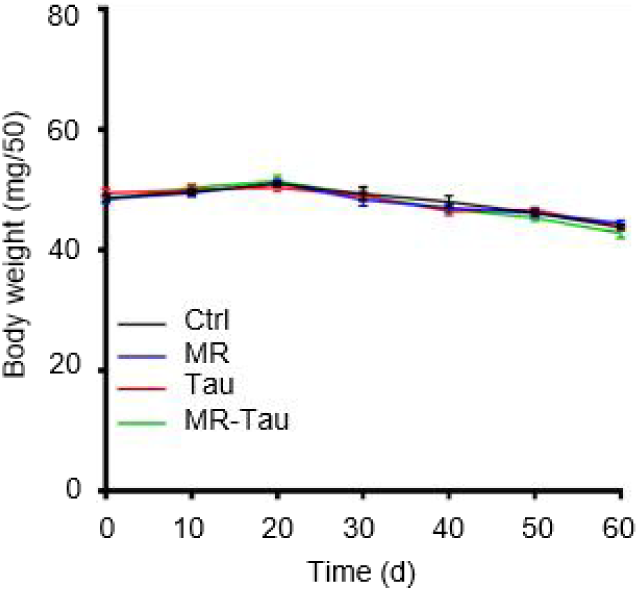
MR and Tau diet have no effect on body weight gain. Body weight measurements across diets. n=150 (male n = 100, female n = 50). All experiments were independently repeated three times. Data are presented as mean ± s.e.m.

**Extended Data Figure 3.**
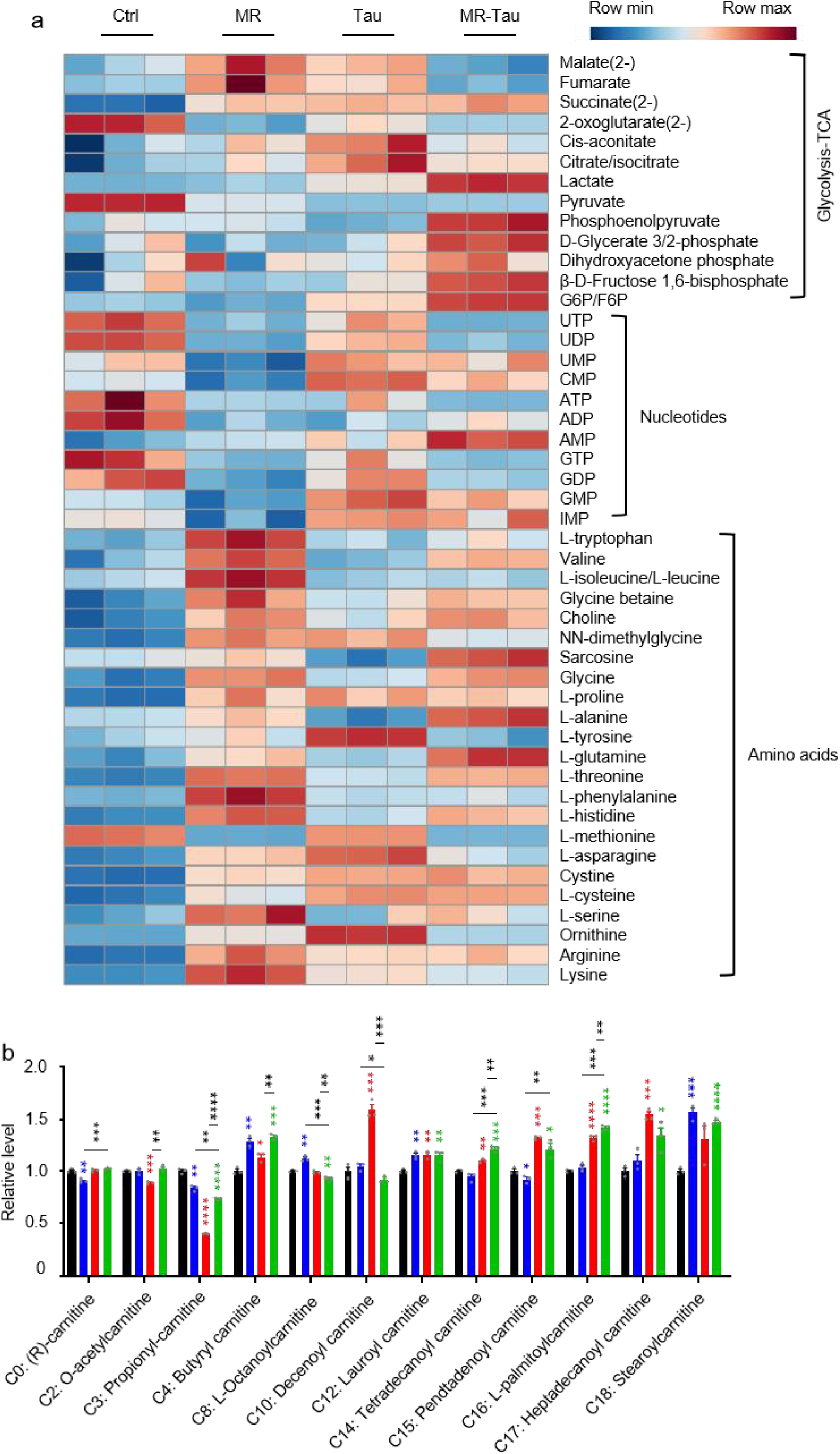
Precision diet modulates carnitine and central carbon metabolism. (a) Heatmap. (b) Carnitine levels. n=10 flies. All experiments were independently repeated three times. The blue, red, and green asterisks indicate the results of MR, Tau, and MR-Tau compared with the control, respectively. Data are presented as mean ± s.e.m. \**P*<0.05, \*\**P*<0.01, \*\*\**P*<0.001, \*\*\*\**P*<0.0001, *P* values were obtained by unpaired, two-tailed *t*-tests unless otherwise specified.

**Extended Data Figure. 4.**
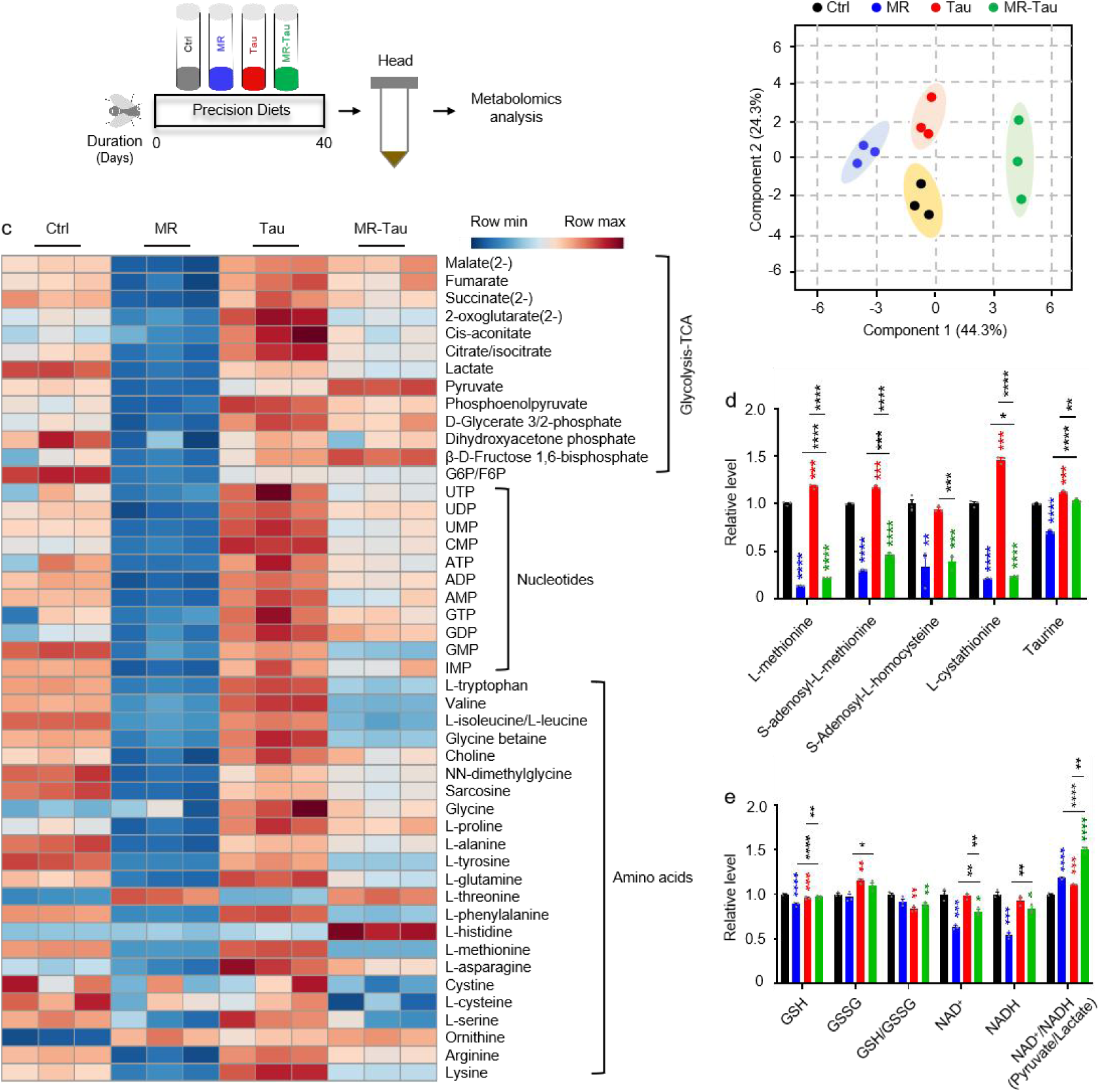
Precision diet reshapes head metabolic landscape. (a) Experimental design. (b) Principal component analysis (PCA). (c) Heatmap. (d) Methionine-related and taurine metabolites. (e) Redox balance. n=30. All experiments were independently repeated three times. The blue, red, and green asterisks indicate the results of MR, Tau, and MR-Tau compared with the control, respectively. Data are presented as mean ± s.e.m. \**P*<0.05, \*\**P*<0.01, \*\*\**P*<0.001, \*\*\*\**P*<0.0001, *P* values were obtained by unpaired, two-tailed *t*-tests unless otherwise specified.

**Extended Data Figure 5.**
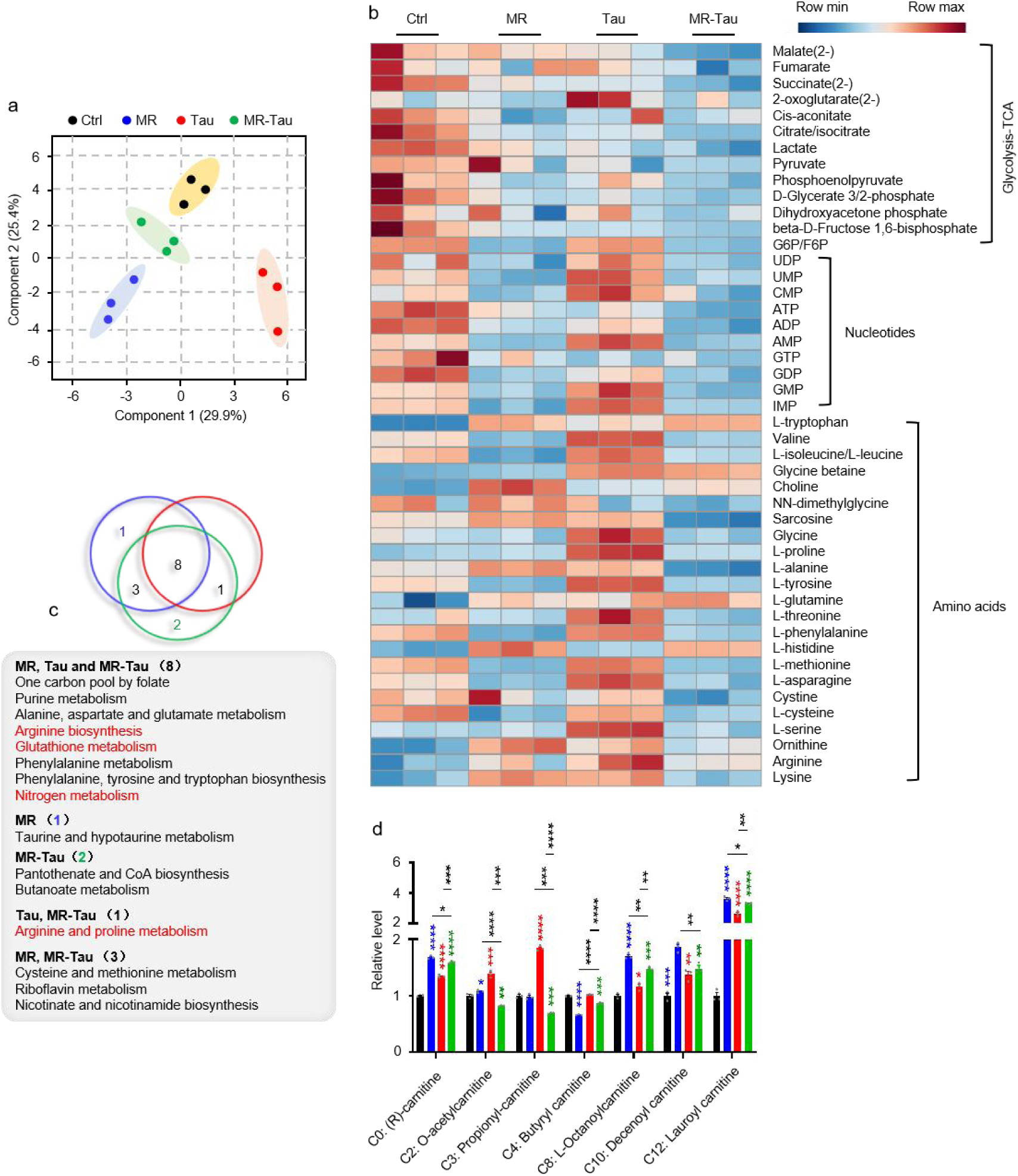
Precision nutrition regulates gut and microbial metabolic homeostasis. (a) PCA. (b) Pathway enrichment analysis. (c) Heatmap. (d) Carnitine levels. n=30. All experiments were independently repeated three times.Data are presented as mean ± s.e.m.

**Extended Data Figure 6.**
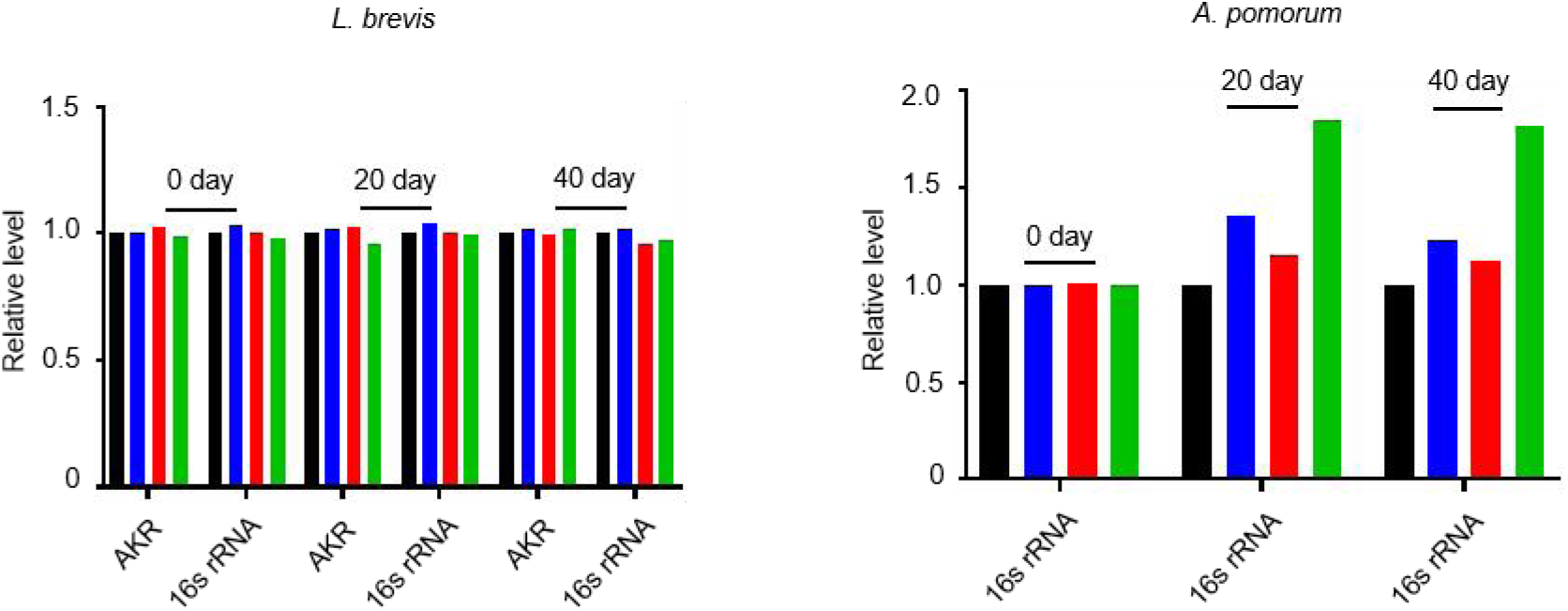
Precision nutrition regulates gut and microbial metabolic homeostasis. (a) Effect of combined intervention on *L. brevis* abundance. (b) Effect of combined intervention on *A. pomorum* abundance. AKR, aldo/keto reductase; *L. brevis-*specific gene. n=30. All reactions were run in five technical replicates. Data are presented as mean ± s.e.m.

**Extended Data Figure 7.**
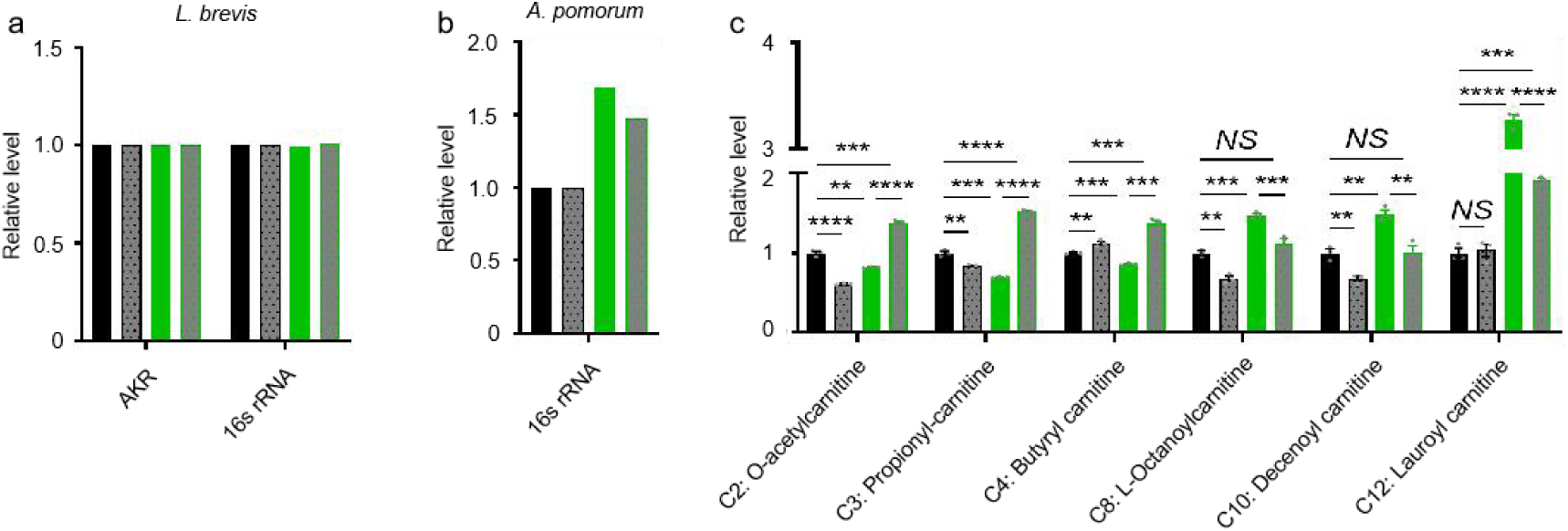
Exercise synergizes with precision nutrition to remodel gut metabolism. (a) Effect of combined intervention on the abundance of *L. brevis*. (b) Effect of combined intervention on the abundance of *A. pomorum*. (c) Carnitine levels. n=30. All experiments were independently repeated five times (a, b) and three times (c). Data are presented as mean ± s.e.m. *NS*, not significant *P*>0.05; \*\**P*<0.01, \*\*\**P*<0.001, \*\*\*\**P*<0.0001, *P* values were obtained by unpaired, two-tailed *t*-tests unless otherwise specified.

**Extended Data Figure 8.**
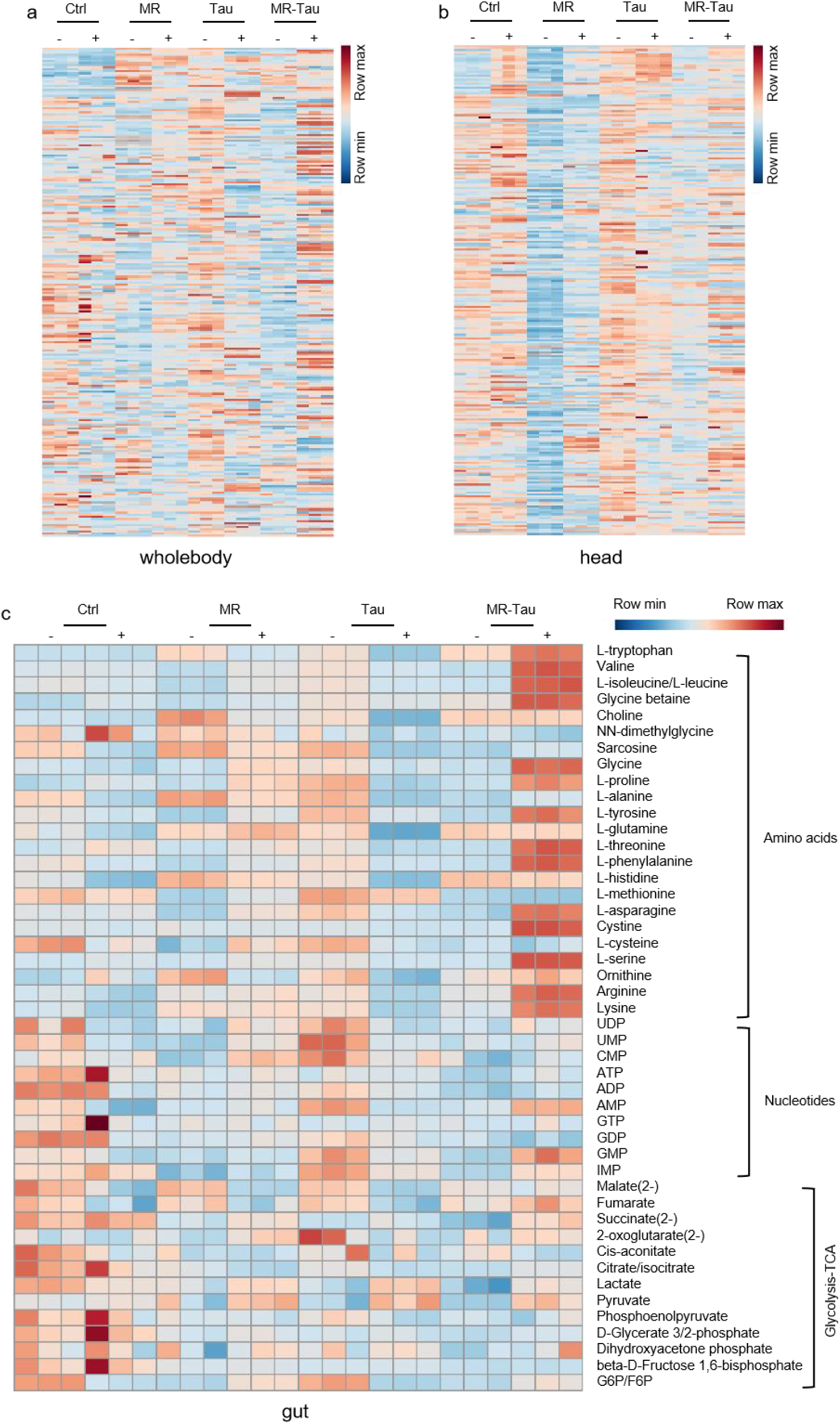
Exercise synergizes with precision nutrition to remodel metabolism. (a) Metabolite enrichment analysis in whole-body (n=10). (b) Metabolite enrichment analysis in head (n=30). (c) Metabolite enrichment analysis in gut (n=30). All experiments were independently repeated three times. Data are presented as mean ± s.e.m.

## Notes

### Competing Interest Statement

The authors have declared no competing interest.

